# METTL7A improves bovine IVF embryo competence by attenuating oxidative stress

**DOI:** 10.1101/2024.12.17.628915

**Authors:** Linkai Zhu, Hao Ming, Giovanna N. Scatolin, Andrew Xiao, Zongliang Jiang

## Abstract

In vitro fertilization (IVF) is a widely used assisted reproductive technology to achieve a successful pregnancy. However, the acquisition of oxidative stress in embryo in vitro culture impairs its competence. Here, we demonstrated that a nuclear coding gene, methyltransferase-like protein 7A (METTL7A), improves the developmental potential of bovine embryos. We found that exogenous METTL7A modulates expression of genes involved in embryonic cell mitochondrial pathways and promotes trophectoderm development. Surprisingly, we discovered that METTL7A alleviates mitochondrial stress and DNA damage and promotes cell cycle progression during embryo cleavage. In summary, we have identified a novel mitochondria stress eliminating mechanism regulated by METTL7A that occurs during the acquisition of oxidative stress in embryo in vitro culture. This discovery lays the groundwork for the development of METTL7A as a promising therapeutic target for IVF embryo competence.

**Summary statement (Graphic abstract):** We describe a molecule acts in the pre-implantation period to attenuate oxidative stress that enhances embryo development to the blastocyst stage and subsequent pregnancy in cattle.

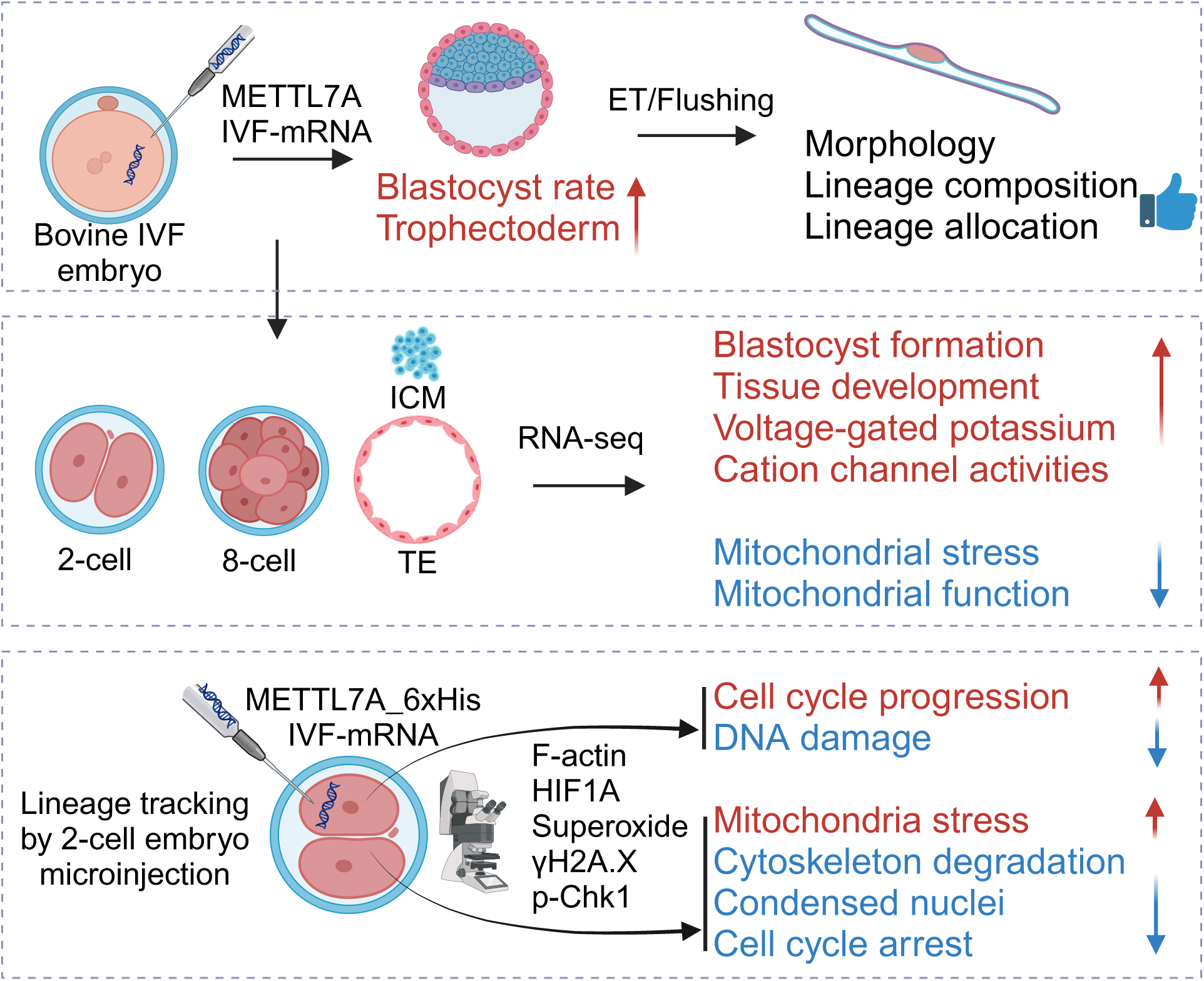

## Introduction

Embryo in vitro production (IVP) technology has been widely used to treat human infertility and improve the reproduction efficiency of agricultural species, such as cattle. The number of IVP embryos transferred has steadily increased over the years globally both from humans [1] and domestic species [2]. However, the competence of IVP embryos to establish pregnancy is much lower than the embryos produced in vivo. It is believed these complications associated with IVF embryos are induced by the environmental stressors accumulated during in vitro embryo culture [3, 4].

Gametes and embryos are exposed to high levels of oxidative stress in in vitro culture conditions [5, 6]. Oxygen (O_2_) tension is one of the major sources that lead to oxidative stress [7]. The atmospheric concentration of O_2_ (20%) used in the embryo culture system is considerably greater than the oxygen tension in the oviduct and uterus of mammals [8]. Studies across multiple species have shown improved in vitro embryo development when oxygen levels are reduced from 20% to 5% [9]. This elevated O_2_ level during embryonic development can influence gene expression, metabolism, and the activity of important epigenetic enzymes.

Another major contributor of oxidative stress is reactive oxygen species (ROS), which are by- products of oxidative phosphorylation within the mitochondria [10]. Under physiological conditions, ROS level is closely monitored and controlled by antioxidants [11]. However, mitochondrial dysfunction and impaired antioxidant defense system can lead to the generation of excess ROS, resulting in delayed development, DNA damage, apoptosis or lipid peroxidation [12].

To mitigate the detrimental effect of oxidative stress on IVP embryo culture to promote embryo competence, multiple approaches have been explored in addition to the reduced oxygen tension, including co-culture with cumulus cells as ROS scavenger [13, 14], supplement of exogenous antioxidants to reduce ROS such as anethole [15], beta-mercaptoethanol [16, 17], imperatorin [18], N-(2-mercaptopropionyl)-glycine [19], and dihydromyricetin [20]. Boosting the endogenous antioxidants such as reduced GSH has also been tested [21]. GSH is the main non-protein sulfhydryl compound in mammalian cells with ability to protect cells from oxidative stress [22].

The synthesis of GSH is dependent on the availability of cysteine, of which intracellular cysteine can be acquired from methionine metabolism [23]. During this process, methionine is first metabolized into S-adenosyl-L-methionine (SAM), which can donate its methyl group to a wide range of molecules catalyzed by methyltransferases and yield S-adenosyl-homocysteine (SAH). SAH undergoes a series of hydrolysis reactions to release free cysteine for GSH synthesis [23].

Recently, a nuclear coding gene, methyltransferase-like protein 7A *(METTL7A),* has been characterized as an endoplasmic reticulum (ER) transmembrane protein involving in lipid droplet formation [24, 25], and it possesses SAM-dependent thiol methyltransferase activity, also named as *TMT1A* [26], thus, it may potentially regulate GSH synthesis. So far, limited studies have shown METTL7A is associated with the successful stem cell reprogramming trajectory [27], and can promote cell survival and modulate metabolic activities [28]. However, the biological function of METTL7A is largely unknown and the specific role of METTL7A during early embryonic development remains unexplored. The aim of this study has been to test the hypothesis that METTL7A alleviates oxidative stress and improves the embryo competence using bovine IVF embryos.

## Materials and Methods

### Animal care and use

Bovine peri-implantation embryos were collected from non-lactating, 3-year-old crossbreed (Bos taurus x Bos indicus) cows. The animal experiments were conducted under animal use protocols (202300000191) approved by the Institutional Animal Care and Use Committee of the University of Florida. All cows were housed in open pasture, and under constant care of the farm staff.

### Bovine oocytes and in vitro embryo production

Germinal vesicle stage oocytes (GV oocytes) were collected as cumulus-oocyte complexes (COCs) aspirated from slaughterhouse ovaries. In vitro maturation was conducted using BO-IVM medium (IVF Bioscience, Falmouth, UK) for 22-23 hours at 38.5°C with 6% CO_2_ to collect MII oocytes. Cryopreserved semen from a Holstein bull with proven fertility was prepared with BO-SemenPrep medium (IVF Bioscience, Falmouth, UK) and added to drops containing COCs with a final concentration of 2 x 10^6^ spermatozoa/ml for in vitro fertilization. Gametes were co-incubated under 38.5 °C and 6% CO_2_. After 10 hours (microinjected embryo experiments) or 16 hours (non-microinjected embryo experiments) in BO-IVF medium (IVF Biosciences, Falmouth, UK), IVF embryos were denuded from cumulus cells by vortexing for 5 min in BO-Wash medium (IVF Bioscience, Falmouth, UK) and cultured up to 7.5 d in BO-IVC medium (IVF Biosciences, Falmouth, UK) at 38.5 °C, 6% CO_2_, and 6% O_2_. Different developmental stage embryos were then evaluated under light microscopy following embryo grade standards of the International Embryo Technology Society. Cleavage rate and blastocyst rate, defined as percentage of cleavage embryos and blastocysts over presumptive zygotes, were measured at embryonic day 3.5 (E3.5) and 7.5 (E7.5), respectively.

### In vitro transcription of *METTL7A*

Total RNA was extracted from a pool of 50 bovine IVF embryos (E7.5), embryonic stem cells, and trophoblast stem cells [29, 30], followed by first strand cDNA synthesis using SuperScript™ IV VILO™ Master Mix (Thermo Fisher Scientific, Waltham, MA). Primers were designed based on the current genome annotation (ARS-UCD2.0; National Center for Biotechnology Information) to include 5’- and 3’- UTR regions of METTL7A (Table S1). PCR was conducted using Q5 Hot Start High-Fidelity 2X Master Mix (New England Biolabs, Ipswich, MA) with an initial denaturation step at 98°C for 30 seconds followed by 30 cycles at 98°C for 10 seconds, annealing at 58°C for 30 seconds and extension at 72 °C for 30 seconds and a final extension at 72 °C for 2 minutes. The purified PCR products were served as DNA template for in vitro transcription using HiScribe® T7 ARCA mRNA Kit (New England Biolabs, Ipswich, MA) with tailing following manufacture’s instruction. The yield and integrity of resulting mRNA were assessed using Qubit 4 (Thermo Fisher Scientific, Waltham, MA) and Tapestation 4150 (Agilent Technologies, Santa Clara, CA).

To generate *METTL7A-6xHis* mRNA, two pairs of primers (Table S1) were designed to produce PCR fragments with an overlapping region which contains 6xHis sequence before the stop codon of *METTL7A*. The two fragments were then assembled using NEBuilder® HiFi DNA Assembly Master Mix (New England Biolabs, Ipswich, MA), followed by in vitro transcription using using HiScribe® T7 ARCA mRNA Kit (New England Biolabs, Ipswich, MA). To overexpress METTL7A, in vitro transcribed mRNAs (IVT-mRNAs) were microinjected into presumptive zygotes or one blastomere of the 2-cell embryo at final concentration of 10 ng/ul.

Approximately 1 pl of solution was injected. In each replicate, approximately 30 presumptive zygotes were injected in one section and cultured together in 50 µl BO-IVC medium. Embryo development rates were recorded from different batches/days of experiments. Data points were excluded only when blastocyst rate from IVF control dropped below 30%, particularly during hot seasons when heat stress may affect oocyte quality [31]. Nuclease-free Tris-EDTA buffer (TEKNOVA T0223, Hollister, CA), which was used to resuspend IVT-mRNAs, was injected in the vehicle control group.

### Immunofluorescence and Data Analysis

Bovine embryos and embryonic cells from different batches/days of experiments were fixed in freshly made 4% paraformaldehyde (Electron Microscopy Science, Hatfield, PA) at room temperature for 15 minutes followed by permeabilization in 1% (v/v) Triton X-100 (Sigma, Burlington, MA) for 20 minutes and blocking at room temperature for 1 hour in 0.1% (v/v) Triton X-100, 0.1M glycine, 2.5% (w/v) BSA (Sigma, Burlington, MA) and 2.5% (v/v) corresponding serum from the host where the secondary antibodies were derived. Samples were then incubated with primary antibodies (Table S1) at 4°C overnight. After three washes in 0.1% (v/v) Triton X-100 and 0.1% (w/v) polyvinylpyrrolidone (PVP; Sigma, Burlington, MA) in Dulbecco’s phosphate buffered saline (DPBS; Thermo Fisher Scientific, Waltham, MA), secondary antibodies (Table S1) were added and incubated at room temperature for 1 hour followed by three washes and mounting on the slide. Confocal images were taken with Olympus IX81-DSU (Cytometry Core Facility, University of Florida, RRID:SCR 019119) and analyzed with ImageJ (V1.53, National Institutes of Health).

### Embryo Transfer

After zygotic injection, ten E6 morulae were transferred to each recipient cows (n = 2) following synchronization with initial intramuscular injection of gonadotropin-releasing hormone (Fertagyl; Merck, Rahway, NJ), standard 7-day vaginal controlled internal drug release (EAZI-BREED CIDR; Zoetis, Parsippany-Troy Hills, NJ) of progesterone, one does of prostaglandin (Lutalyse; Zoetis, Parsippany-Troy Hills, NJ) upon CIDR removal and another dose of gonadotropin-releasing hormone 48 hours after CIDR removal. Heat detection was determined by scratch of ESTROTECT patches (ABS GLOBAL, DeForest, WI). A cohort of ten morulae from control or treatment group were loaded into 0.5 ml straws in prewarmed Holding Medium (ABT 360, Pullman, WA) and transferred non-surgically to the uterine horn ipsilateral to corpus luteum as detected by transrectal ultrasound. Embryos were recovered by standard non-surgical flushing with lactated ringer’s solution (ICU Medical, San Clemente, CA) supplemented with 1% (v/v) fetal bovine serum on embryonic day 12. After flushing, all surrogate cows were given one dose of 5 ml prostaglandin.

### Interferon-Tau Assay

Blood samples from recipients were collected from the coccygeal vein using serum separator tubes on the day of flushing, and immediately stored in refrigerator before centrifugation for 15 minutes at 1000 x g. Serum interferon tau (IFNτ) level was measured with Bovine Interferon-Tau ELISA Kit (CUSABIO, Houston, TX) per manufacture’s instruction. Briefly, 100 µl standard or sample were added to each well of 96-well plate provided in the kit and incubated for 2 hours at 37 °C. Liquid was withdrew and 100 µl biotin-antibody was added to each well, followed by 1 hour incubation at 37 °C. The solution was discarded, and the wells were washed three times with 200 µl Wash Buffer. To remove any remaining Wash Buffer in the wells, the plate was inverted and placed on clean paper towel for 1 minute. 100 µl HRP-avidin was then added to each well and incubated for 1 hour at 37 °C followed by five times of washes. For signal detection, 90 µl TMB Substrate was added and incubated for 20 minutes at 37 °C avoiding light. After incubation, 50 µl Stop Solution was added to each well while gently shaking the plate to ensure thorough mixing. The plate was measured using a colorimetric microplate reader set to 450 nm.

### Separation of TE and ICM

To profile the differential transcriptome during first lineage specification, blastocysts with zona pellucida at embryonic day 7.5 were used for TE/ICM dissociation following a previously published protocol (26). Briefly, 0.25% trypsin (Thermo Fisher Scientific, Waltham, MA) was continuously injected into the blastocysts until a small mass of cells was slowly washed out from the zona pellucida. The cell masses were washed three times with 0.1% PVP and immediately transferred to -80°C until further use or fixed in freshly made 4% paraformaldehyde followed by staining.

### RNA sequencing analysis

Five 2- or 8-cell embryos were pooled in each replicate and TE and ICM cell clumps from 5 blastocysts were pooled after separation for RNA-seq library preparation. Embryos and cells were used directly for library preparation without RNA extraction following manufacturers’ instructions. Briefly, SMART-Seq v4 Ultra Low Input RNA kit (Takara, Mountain View, CA) was used for cDNA synthesis and amplification. Library preparation was conducted using Nextera XT DNA Library Prep Kit (Illumina, San Diego, CA). The libraries were subject to size selection with 0.6x AMPure XP bead wash (Beckman Coulter, Indianapolis, IN). The concentration of RNA-seq libraries was determined with a Qubit high sensitivity dsDNA HS assay kit (Thermo Fisher Scientific, Waltham, MA). Pooled indexed libraries were then sequenced on the Illumina NovaSeq 6000 platform with 150-bp paired-end reads.

Multiplexed sequencing reads that passed filters were trimmed to remove low-quality reads and adaptors by Trim Galore (version 0.6.7). The quality of reads after filtering was assessed by FastQC, followed by alignment to the bovine reference genome by HISAT2 (version 2.2.1) with default parameters. The output SAM files were converted to BAM files and sorted using SAMtools6 (version 1.14). Read counts of all samples were quantified using featureCounts (version 2.0.1) with the bovine genome as a reference and were adjusted to provide counts per million (CPM) mapped reads. Pearson correlation and Principal Component analysis were performed with R (Version 4.4.1). Differentially expressed genes were identified using edgeR (version 4.2.1) in R. Genes were considered differentially expressed when they provided a false discovery rate (FDR) of <0.05 and |log_2_FC| > 1. Bioconductor package ClusterProfiler (version 4.12.1) was used to reveal the Gene Ontology (GO) and KEGG pathways in R.

### Western Blot and Data Analysis

Ten E7.5 blastocysts were washed three times in 0.1% (w/v) PVP and pooled with approximately 5 µl medium carryover in each replicate. Samples were first heated with 5 µl 2xSDS gel-loading buffer at 95 °C for 5 minutes followed by a quick spin down, and then loaded onto 10% Tris-Glycine Mini Protein Gels (Thermo Fisher Scientific, Waltham, MA). Western blot electrophoresis was conducted at 100 volts for 2 hours. Proteins were transferred from gel to PVDF membrane with an iBlot 3 Western Blot Transfer Device (Thermo Fisher Scientific, Waltham, MA). After transfer, the membrane was washed with 25 ml Tris buffered saline (TBS) for 5 minutes at room temperature followed by blocking with 2.5% (w/v) BSA and 2.5% (v/v) corresponding serum, from the host where the secondary antibodies were derived, for 1 hour at room temperature. The membrane was washed three times for 5 minutes each with 15 ml of TBST (0.1% Tween-20 in TBS). Primary antibodies were added in 10 ml dilution buffer (5% w/v BSA in TBST) and incubated with membrane with gentle agitation overnight at 4°C followed by three times of washes with TBST. Secondary antibodies were added in 10 ml of blocking buffer and incubated with membrane with gentle agitation for 1 hour at room temperature. Membrane was washed three times before proceeded with signal detection. Pierce™ ECL Western Blotting Substrate (Thermo Fisher Scientific, Waltham, MA) was added to the membrane and incubated for 1 minute. Excessive solution was removed before imaging using iBright CL1500 System (Thermo Fisher Scientific, Waltham, MA).

### Superoxide Assay

MitoSOX Green (MSG, Thermo Fisher Scientific, Waltham, MA) reagent stock was prepared by dissolving the contents of the vial in 10 µl of anhydrous DMF, which is stable for one day. To make a working solution, 10 µl of 1mM stock solution was added to HEPES-TALP. 200 µl of working solution was added to one well of µ-Slide (Ibidi, Fitchburg, WI). Live embryos were taken out from culture and washed quickly with HEPES-TALP followed by incubation in MSG working solution for 30 minutes at 38.5 °C, 6% CO_2._ After incubation, embryos were washed three times with warm buffer and confocal images were taken within 2 hours of staining.

Fluorescence intensity was analyzed with ImageJ and Prism 9 (GraphPad, La Jolla, CA). A two-tailed student’s t-test was used for statistical analysis.

### GSH Assay

To measure the level of glutathione in reduced form (GSH), ten 8-cell embryos or five E7.5 blastocysts were pooled in each replicate after washing briefly in 0.1% (w/v) PVP/PBS. The level of reduced GSH was quantified indirectly by subtracting oxidized GSH (GSSG) from total GSH using commercial kit GSH/GSSG-Glo™ Assay (Promega, Madison, WI) following manufacturer’s instructions. Relative luminescence over no cell control was analyzed with two-tailed student’s t-test in Prism 9.

## Results

### Exogenous METTL7A improves the developmental potential of bovine IVP embryos

By mining of genome-wide transcriptional and translational datasets [32], we found that METTL7A is barely expressed and translated across bovine oocytes and pre-implantation embryos derived in vitro (Figure S1), suggesting that METTL7A is dispensable for bovine pre- implantation development. To test the biological function of METTL7A during embryogenesis, we microinjected exogenous METTL7A mRNA into zygotes (Table S1) and evaluated the effect of overexpression (OE) of METTL7A during bovine pre-implantation development. While there is no difference in cleavage rate between METTL7A ^OE^ and the control group (79.29% vs. 81.42%; N= 980, n =29 for the treatment group; *p* = 0.4503), a 14.32% increase in blastocyst formation rate was observed in METTL7A ^OE^ compared to controls (54.96% vs. 40.64%; N= 335, n =11 for the treatment group; *p* = 0.0102) (Figure 1A, B), indicating a beneficial role of METTL7A for bovine pre-implantation development. Additionally, METTL7A^OE^ blastocysts had a normal differentiation into inner cell mass (ICM) and trophectoderm (TE), as assessed by immunostaining analysis of SOX2 and CDX2 (Figure 1C, D), respectively. Notably, the number of TE cells was significant higher in the METTL7A ^OE^ blastocysts compared to the control group (134 vs. 87; n = 5; *p* = 0.0198), while ICM cell number was not different (29 vs. 27.6; n = 5; *p* = 0.5548), resulting in higher TE/ICM ratio in METTL7A ^OE^ blastocysts compared to control (4.81 vs. 3.16, *p* = 0.0699) (Figure 1E-G).

**Figure 1.**
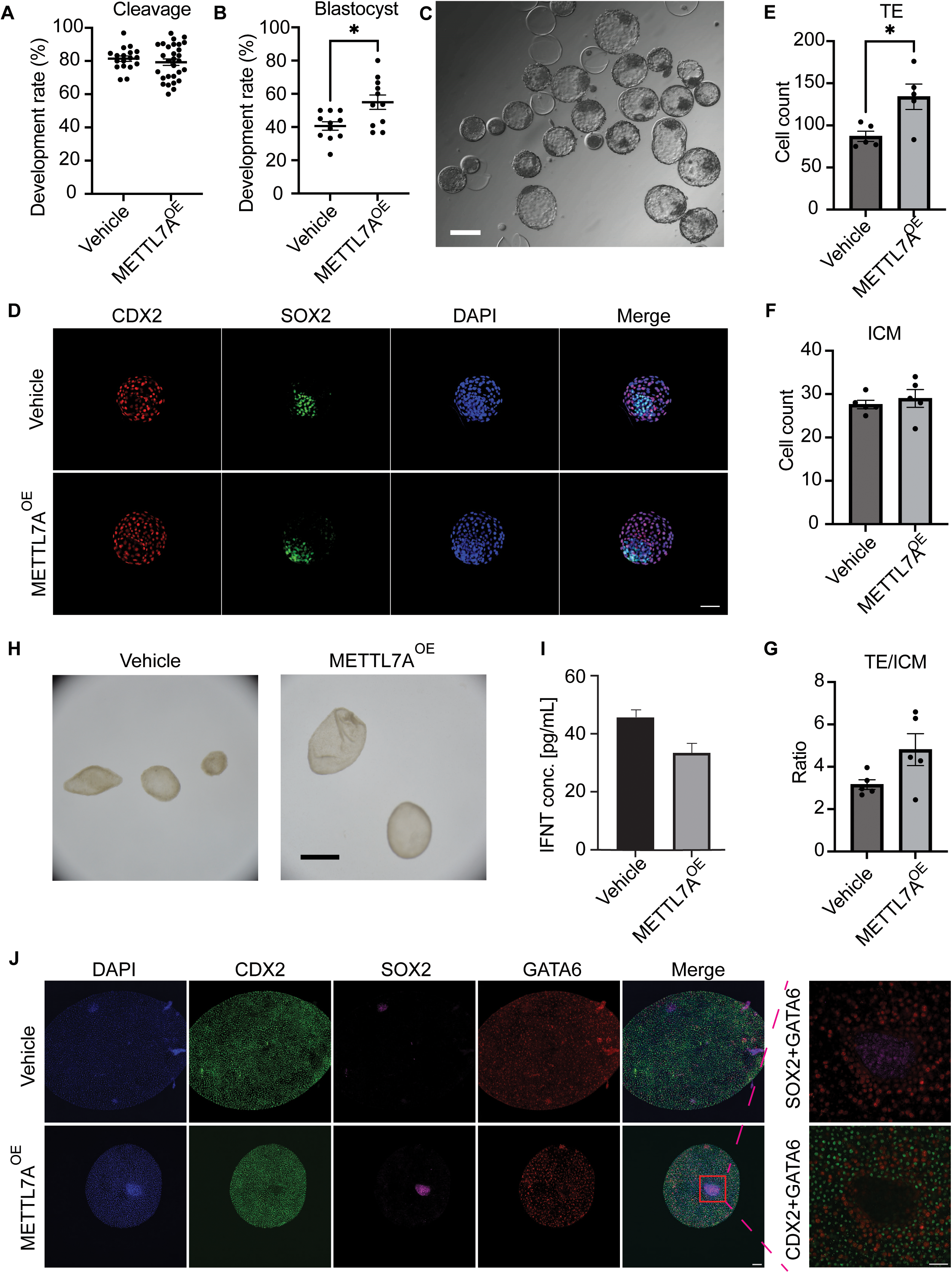
Exogenous METTL7A improves the developmental potential of bovine IVP embryos. **A**. The cleavage OF METTL7A ^OE^ embryos (n = 29) compared to the vehicle control (n = 18). **B**. The blastocyst rate of METTL7A ^OE^ embryos (n = 11) compared to the control (n = 11). **C**. A representative image of bovine METTL7A ^OE^ embryos, scale bar = 100 µm. **D**. Immunostaining analysis of CDX2 (trophectoderm, TE marker) and SOX2 (inner cell mass, ICM marker) in METTL7A ^OE^ embryos compared to the control, scale bar = 50µm. **E**. The total TE cell number counts between METTL7A ^OE^ blastocysts (n = 5) compared to the control (n = 5). **F**. The total ICM cell number counts between METTL7A ^OE^ blastocysts (n = 5) compared to the control (n = 5). **G**. The TE/ICM cell number ratio between METTL7A ^OE^ blastocysts compared to the control. **H**. A representative bright field image of day 12 (E12) elongated embryos from METTL7A ^OE^ and control embryo transfer, scale bar = 500 µm. **I**. The serum INF-tau levels of surrogate cows with METTL7A ^OE^ embryo transfer (n = 2) compared to the control (n = 2) on the day of flushing. **J**. Immunostaining analysis of CDX2 (TE marker), SOX2 (embryonic disc marker), and GATA6 (hypoblast marker) in E12 embryos flushed out from the METTL7A ^OE^ and control embryo transfer, scale bar = 50µm.

To further determine the viability of METTL7A ^OE^ embryos and if they can establish successful pregnancy, we transferred either METTL7A ^OE^ or IVF embryos at morula stage to recipient cows and flushed them out on embryonic day 12 (E12) for analysis. We found METTL7A ^OE^ embryos displayed normal morphology and lineage differentiation similar to the control group (Figure 1H, J), and initiated maternal recognition of pregnancy as indicated by a comparable serum INF-tau level in the surrogate mothers as IVF embryo transfers (n = 2, Figure 1I).

These results demonstrated that METTL7A promotes the developmental potential of bovine pre- implantation embryos by facilitating TE lineage development, and that bovine METTL7A ^OE^ embryos produce normal pregnancy through conceptus elongation following embryo transfer to recipients. Overall, these results highlight that METTL7A molecule constitutes a promising pharmaceutical target for improving IVF embryo competence.

### Exogenous METTL7A modulates expression of genes involved in mitochondrial functions during bovine pre-implantation development

To understand METTL7A function on gene expression of embryos and embryonic (ICM and TE) lineages, we performed RNA sequencing (RNA-seq) analysis on METTL7A ^OE^ and control embryos at 2-, 8-cell and blastocyst stage. ICM and TE were separated by micromanipulation procedures, which were confirmed by immunostaining analysis of lineage markers SOX2 and CDX2, respectively (Figure S2A, B). Pearson correlation and principal component analysis of transcriptomic data indicated consistent values between biological replicates across developmental stage (Figure 2A, B). While the transcriptomes of both METTL7A ^OE^ and control embryos were distinct across developmental stages, they appeared to cluster together within the same stage with more notable differences in 2 and 8-cell embryos than blastocysts (ICM and TE) (Figure 2A, B).

**Figure 2.**
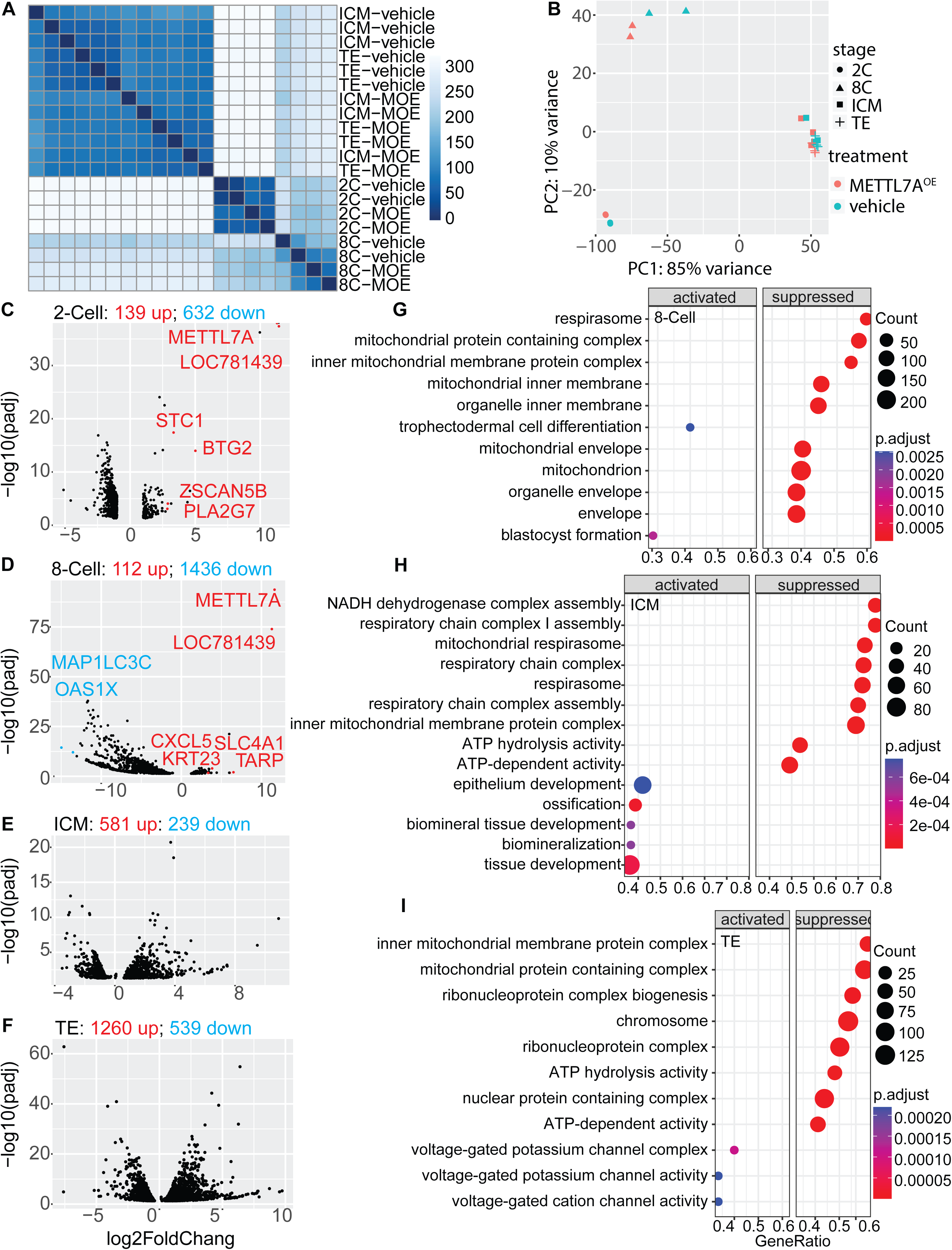
The transcriptomic analysis of METTL7A ^OE^ embryos at 2-, 8-cell, and ICM/TE compared to control. **A**. Heatmap of the samples from the same stages of bovine embryos from the METTL7A ^OE^ and control group. **B**. Principal component analysis (PCA) of the transcriptomes of METTL7A ^OE^ and control embryos at different developmental stages. **C-F** Volcano plots showing the number of up- or down-regulated genes in METTL7A ^OE^ embryos compared to control at 2-cell (C), 8-cell (D), ICM (E) and TE (F) stages (FDR < 0.05, |log_2_FC| > 1). The most significant up-regulated genes in METTL7A ^OE^ embryos compared to the control are highlighted. (**G-I**) The top GO terms of up- and down-regulated genes METTL7A ^OE^ embryos compared to the control at of 8-cell (G), ICM (H) and TE (I) stages.

In 2-cell embryos, we found 139 and 632 genes to be up- and down-regulated (FDR P value < 0.05, |log_2_FC| >1) in METTL7A ^OE^ compared to control embryos, respectively (Figure 2C). The most up-regulated genes include *METTL7A*, *LOC781439, BTG2, STC1, ZSCAN5B*, and *PLA2G7* (Figure 2C, Table S2). Of note, *METTL7A* and *LOC781439 (*a pseudogene with truncated sequence of *METTL7A)* were the most regulated genes *with* log_2_FC > 11 (Table S2), confirming the overexpression of METTL7A.

At the 8-cell stage, 112 and 1,436 genes were up- and down-regulated in METTL7A ^OE^ compared to control embryos, respectively (Figure 2D, Table S3). Similarly, *METT7A* and *LOC781439* remained the top two up-regulated genes with log_2_FC > 11 (Table S3), indicating *METTL7A* overexpression pertains to the 8-cell stage. Other top up-regulated genes included *KRT23, TARP, CXCL5, and SLC4A10* (Table S3), with known biological roles in promoting proliferation [33], DNA damage response [34], tumor progression [35], and pH balance [36], respectively. Compared to 2-cell stage, there were significant more down-regulated genes in METTL7A ^OE^ embryos at the 8-cell stage. Only one gene had log_2_FC < -5 at the 2-cell stage while 401 genes showed log_2_FC < -5 at the 8-cell stage (Figure 2C, D, Table S2, S3). Most of the top down-regulated genes in 8-cell embryos (Table S3) were associated with various stress responses, such as *MAP1LC3C* (as known as *LC3C*) and *OAS1X* that are responsible for antibacterial and antiviral response [37, 38].

At the blastocyst stage, we observed a larger number of genes differentially expressed in TE than ICM (up-regulated: 1,260 vs. 581; down-regulated: 539 vs. 239) associated with METTL7A overexpression (Figure 2E, F; Table S4, S5). However, METTL7A was no longer up-regulated in the blastocysts (both TE and ICM) (Table S4, S5), indicating that the observed transcriptomic changes were not directly caused by overexpression of *METTL7A* but rather from an altered gene expression cascade induced from cleavage stages.

Gene ontology (GO) analysis indicated overexpression of METTL7A suppressed genes involved in mitochondrial stress and functions among 8-cell and blastocyst (ICM and TE) stage embryos (Figure 2G-I). On the contrary, the up-regulated genes by overexpression of METTL7A were involved in blastocyst formation at 8-cell stage, tissue development in ICM cells, and voltage-gated potassium and cation channel activities in TE cells, respectively (Figure 2G-I). Given that stress responses demand high energy consumption provided by mitochondria [39], the RNA-seq results suggested that overexpression of METTL7A shifts the paradigm of energy expenditure to favor bovine embryonic development.

Additionally, genes were found to be precisely modulated in the presence of METTL7A by comparing datasets between stages. For example, there is an earlier activation and up- regulation of HAND1 observed at the 2-cell and the 8-cell embryos (Table S2, S3). HAND1 is essential for trophoblast lineage differentiation and development [40]. However, at the blastocyst stage, the expression of HAND1 remain unchanged (Table S4, S5), coinciding with the termination of METT7A overexpression in this stage.

### Exogenous METTL7A reduces mitochondrial stress and decreases superoxide level of bovine pre-implantation embryos

Given mitochondria stress is precisely regulated in embryos and is associated with embryo competence, and that the mitochondria related pathways are among the top regulated among METTL7A ^OE^ embryos at 2-, 8-cell and blastocyst stage, we next sought to determine the effect of exogenous METTL7A on embryonic cell mitochondrial stress. We introduced a 6x His Tag before the stop codon of METTL7A due to the lack of a suitable bovine METTL7A antibody (Figure S2C, Table S1) and established a lineage tracking system by 2-cell embryo microinjection. The intracellular localization of the expressed METTL7A-6xHis fusion protein was evaluated 12 hours after injecting the in vitro transcribed mRNAs into one of the blastomeres at the 2-cell stage (Figure 3A). We confirmed that METTL7A was uniformly distributed into the cytoplasm, with no enrichment in particular organelles (Figure 3B). We found that METTL7A-positive blastomeres progressed through one or two cell cycles within 12 hours, whereas METTL7A-negative blastomeres were arrested with condensed nuclei and degraded cytoskeleton (Figure 3B), indicating an apoptotic cell fate [41]. Moreover, METTL7A-negative blastomeres exhibited activation and nuclear translocation of HIF-1α (Figure 3B), a maker of mitochondrial adaptation to oxidative stress [42]. These results suggested that METTL7A can protect IVF embryos from mitochondrial stress, which was further supported by the presence of attenuated mitochondrial respiratory chain activities in METTL7A ^OE^ embryos (10 blastocysts/replicate, *p* = 0.0142) (Figure 3C-E), and was consistent with down-regulation of mitochondrial pathways in METTL7A ^OE^ embryos (Figure 2G-I).

**Figure 3.**
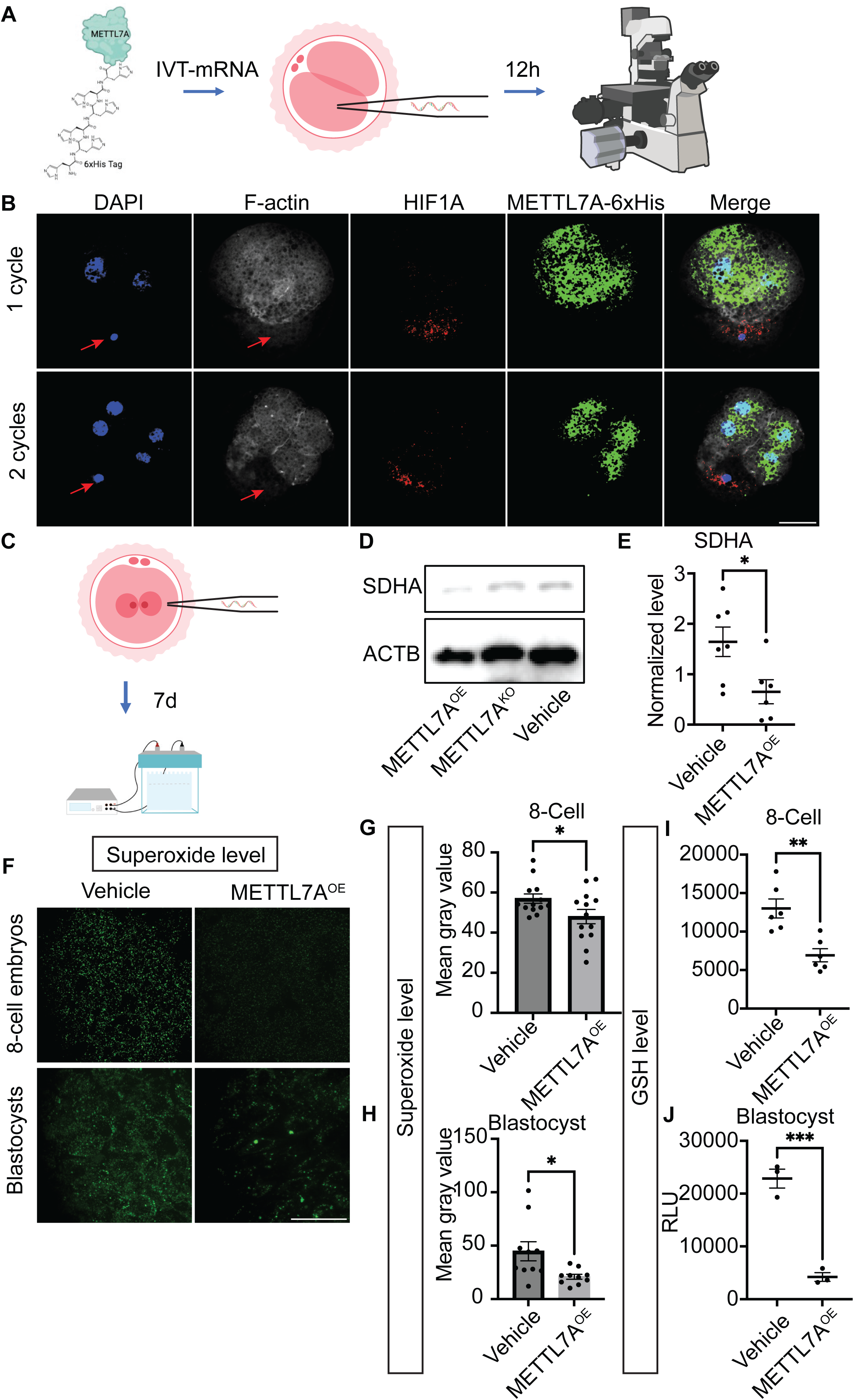
Exogenous METTL7A reduces mitochondrial stress and decreases superoxide level of bovine pre-implantation embryos. **A**. Experimental scheme of injecting METTL7A IVT-RNA into one blastomere of 2-cell embryos. **B**. Immunostaining analysis of F-actin, mitochondrial stress marker (HIF1a), and METTL7A-6xHisTag in METTL7A ^OE^ blastomeres and control. The red arrow points to condensed chromatin and degradation of actin filament, scale bar = 50µm. After 12 hours of injection, METTL7A ^OE^ blastomeres developed through 1 or 2 cell cycles, while non-injected blastomeres were arrested (n = 5, 4/5 or 80%) with described phenotype. **C**. Experimental scheme of zygotic injection and western blot analysis. **D** and **E**. Western blot analysis of Succinate Dehydrogenase Complex Flavoprotein Subunit A (SDHA, a marker for mitochondrial respiratory activity) in METTL7A ^OE^ (n = 6) blastocysts compared to control (n = 7). F. The immunostaining analysis of superoxide level measured by MitoSox green in METTL7A ^OE^ embryos and control at 8-cell (n = 13 embryos for both groups) and blastocyst (n = 10 embryos for both groups) stage, scale bar, 50µm. **G** and **H**. The quantification of superoxide level in METTL7A ^OE^ embryos and control at 8-cell (G) and blastocyst stage (H). **I** and **J**. The GSH level METTL7A ^OE^ embryos and control at 8-cell (I, n = 6 for both groups) and blastocyst stage (J, n = 3 for both groups).

To further delineate the mitochondrial stress relief conferred by METTL7A, we measured a cause of oxidative stress, reactive oxygen species (ROS), in METTL7A ^OE^ and control embryos. As expected, superoxide levels were reduced in METTL7A ^OE^ embryos compared to control (8- cell stage, *p* = 0.0454; blastocyst stage, *p* = 0.0182), as measured by the MitoSox green assay (Figure 3F-H). In concordance with lower superoxide levels, the levels of the intracellular antioxidant glutathione in its reduced form (GSH) also decreased dramatically (8-cell stage, *p* = 0.0023; blastocyst stage, *p* = 0.0007) in METTL7A ^OE^ embryos compared to control (Figure 3I, J), indicating active reduction reactions in METTL7A ^OE^ embryos.

Together, these results indicated that METTL7A alleviates mitochondrial stress and oxidative stress during bovine pre-implantation embryo development.

### Exogenous METTL7A attenuates DNA damage and promotes cell cycle progression

ROS are also genotoxic [43], prompting us to evaluate the effect of overexpression of METTL7A on embryonic cell DNA damage. DNA damage occurred in normal IVP embryos (Figure 4A), consistent with previous findings [44]. Using the same blastomere injection approach, we found METTL7A-negative blastomeres had a higher level of DNA damage particularly at 4-cell stage, as measured by yH2A.X staining (Figure 4A). At the blastocyst stage, coincided with higher blastocyst rate, a lower level of DNA damage was observed in METTL7A ^OE^ embryos compared to control (*p* = 0.0288) (Figure 4B, C). These results demonstrated that METTL7A reduces embryonic cell oxidative stress and DNA damage, thereby promoting the developmental potential of bovine IVP embryos.

**Figure 4.**
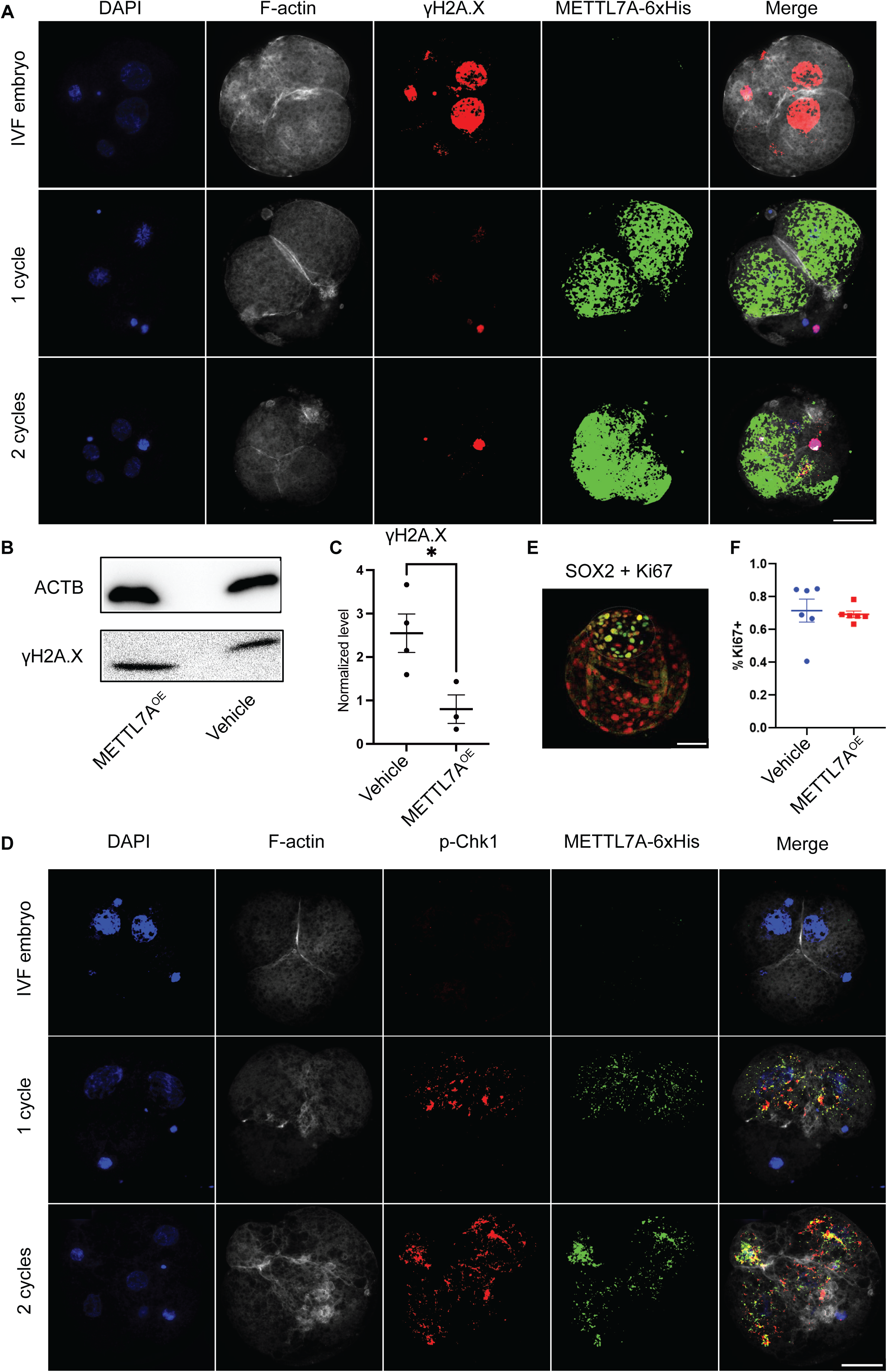
**Exogenous METTL7A attenuates DNA damage and promotes cell cycle progression**. **A**. Immunostaining analysis of F-actin, yH2A.X (DNA damage marker) and METTL7A-6xHisTag in METTL7A ^OE^ blastomeres (n = 5) and control (non-injected blastomeres and IVF embryos: METTL7A ^OE^ negative blastomere), scale bar = 50µm. **B** and **C**. Western blot analysis and quantification of yH2A.X in METTL7A ^OE^ embryos (n = 3) and the vehicle control (n = 4) at blastocyst stage 7 days after zygotic injection. **D**. Immunostaining analysis of p-Chk1 (cell cycle checkpoint marker) in METTL7A ^OE^ and IVF control embryos at the cleavage stage, scale bar = 50µm. **E** and **F**. Immunostaining analysis of the proliferative cells (Ki67^+^) in METTL7A ^OE^ blastocyst and control group. After 12 hours of zygotic injection, METTL7A ^OE^ blastomeres developed through 1 or 2 cell cycles, while non-injected blastomeres were arrested (n = 5).

Given that DNA damage can lead to the delayed cell cycles and impaired embryo development [44], we further analyzed p-Chk1, an essential marker for the DNA damage checkpoint and control of mitotic entry [45], in the METTL7A modulated embryos. We found that p-Chk1 is significantly up-regulated in blastomeres 12 hours post-injection of METTL7A_6xHis Tag (Figure 4D). However, at the blastocyst stage, the percentage of proliferating cells did not differ between METTL7A ^OE^ and control embryos (Figure 4E, F), consistent with our previous observation that a lack of exogenous METTL7A persists into blastocyst stage embryos (Figure 2E, F).

In summary, our results demonstrated that METTL7A ameliorates DNA damage by reducing ROS levels and enhancing DNA damage repair through p-Chk1, which promotes the ‘error-free’ cell cycle progression and pre-implantation embryo development.

## Discussion

In vitro fertilization (IVF) is one of the most used assisted productive technologies in both humans and domestic species. While IVF procedures are considered safe, the in vitro developing embryos are exposure to conditions during culture not normally experienced in vivo, when zygote undergoes extensive epigenetic and metabolic reprogramming, potentially leading to alterations in the embryonic gene expression that may result in adverse outcomes. It has long been established that embryo in vitro culture induces pestilent oxidative stress to embryos [46]. Therefore, there is a critical need to identify the strategy to protect in vitro developing embryos from oxidative stress. Here, we identified a molecule, METTL7A, improves bovine embryo competence by attenuating oxidative stress. Specifically, we found that exogenous METTL7A promotes the developmental potential of bovine pre-implantation embryos by facilitating trophectoderm lineage development and alleviating embryonic cell oxidative stress, and that the resultant embryos produce normal pregnancy through conceptus elongation following embryo transfer to recipients. Our findings could have broader implications for the development of METTL7A molecule for optimal IVF culture conditions.

RNA-seq comparative analysis of METTL7A ^OE^ and control embryos provided interesting observations. For example, among the most regulated genes in 2-cell embryos, BTG2 has been reported to destabilize mRNA [47], while ZSCAN5B is associated to embryonic genome activation through modulating mitotic progression and safeguarding DNA damage response [48–50]. Given the importance of maternal mRNA clearance for embryonic development [51], METTL7A overexpression may facilitate embryonic genome activation via downstream effectors like BTG2 and ZSCAN5B. Additionally, STC1 is hypoxia-responsive and promotes lipid metabolism [52, 53] and its abundance is positively associated with all Ovum Pick Up-In vitro Production (OPU-IVP) scores [54]. Similarly, PLA2G7 (also known as LDL-PLA2) regulates phospholipid catabolism during inflammation and oxidative stress responses [55, 56]. The up- regulation of STC1 and PLA2G7 in METTL7A ^OE^ embryos suggested that METTL7A can alleviate oxidative stress in early bovine embryos.

Mechanistically, our results have demonstrated that this is through regulation of mitochondrial stress and enhancing the utilization of GSH during early embryonic cleavage stages.

Specifically, our study has shown that the superoxide level within the mitochondria was reduced by exogenous METTL7A, indicating an ameliorated mitochondria stress in cleavage embryos, which was further supported by observations of overall lower mitochondrial activities and down- regulation of HIF-1α. Surprisingly, overexpression of METTL7A decreased intracellular GSH level, challenging our speculation that the SAMe-dependent enzymatic activity of METTL7A would contribute to the synthesis of GSH. Also, the GSH assay used in this study was only able to capture intracellular GSH, while the oxidized GSSH diffused out of the cell [57]. Consequently, DNA damage levels were ameliorated, with enhanced cell cycle checkpoint mechanism, therefore ensuring proper cell cycle progression and development into later embryonic stages.

METTL7A has previously been observed to localize to the endoplasmic reticulum and the inner nuclear membrane [24, 25], and thus the biological function of this protein in early embryonic development has not previously been uncovered. Previous work has suggested that METTL7A has methyltransferase activities toward lncRNA and thiol group substrates and could promote stem cell reprogramming [27], and modulate metabolic stress to improve cell survival [28]. Here we demonstrated another function of METTL7A that could shift the paradigm of energy expenditure and ameliorates oxidative stress from mitochondrial metabolism, therefore promoting bovine blastocyst formation in vitro. Our work nevertheless has identified a completely novel, reducing oxidative stress function of this relatively understudied protein.

Future biochemical, genomic and structural work will investigate the direct target molecule network of METTL7A in early embryos.

We have also identified a novel function of exogenous METTL7A in promoting trophectoderm development and modulating the embryonic cell transcriptome associated with mitochondria stress and function for improving embryo survival. Although the direct target molecule network of METTL7A remains unexplored, our findings have shown that several essential transcriptional factors (e.g., HAND1) were precisely modulated in the presence of METTL7A. Meanwhile, the IFN-tau secretion by trophoblast remained unaffected, suggesting normal trophoblast development. Moreover, we couldn’t rule out if the increased trophectoderm cell number by METTL7A overexpression is due to the accelerated timing of blastocyst formation, and if and how reducing oxidative stress promote the progression and timing of blastulation. Future studies to monitor the embryo development in real time with a time-lapse incubator could be of particularly interesting.

Under in vivo conditions, the oviduct and uterus provide abundant antioxidants to tightly control ROS level, creating an optimal environment for embryo growth [58]. However, current culture systems fail to provide a constant antioxidant supply, leading to impaired embryo development [12]. Indeed, by comparing the single blastomere transcriptomic profiles of bovine in vivo and in vitro derived blastocyst, a recent study has shown that there are highly active metabolic and biosynthetic processes, reduced cellular signaling, and reduced transmembrane transport activities in IVP embryos that may lead to reduced developmental potential [59]. Similarly, compared to IVP embryos, the up-regulated genes in vivo embryos involve in regulation of embryonic development and tissue development [59] as we observed in METTL7A ^OE^ embryo transcriptomic datasets. Since our work was all conducted in in vitro conditions, we cannot rule out the possibility that endogenous METTL7A presents in in vivo embryos, thus helps to eliminate any negative oxidative stress to enhance embryo survival. Therefore, future work should carefully access the expression dynamics of METTL7A in in vivo conditions.

Additionally, comprehensive examination of the act of METTL7A in embryos throughput pregnancy and their offspring will pave the utility of METTL7A as a promising pharmaceutical target for IVP embryo viability.

In summary, we have identified a novel mitochondria stress eliminating mechanism regulated by METTL7A that occurs during the acquisition of oxidative stress in embryo in vitro culture. We believe that this work is novel at both the mechanistic and translational levels: METTL7A not only displays a unique oxidative stress relief function, but it also lays the groundwork for the development of strategies that could specifically prevent oxidative stress in IVF. Future work will further investigate the possibility of efficiently deliver METTL7A into IVF embryos and understand its downstream molecular network targets.

## Competing interests

The findings of this study were included in a U.S provisional patent application 63/698,174.

## Author contributions

Conceptualization: Z.J; Methodology: L.Z, H.M, G.S; Validation: L.Z, Z.J; Formal analysis: L.Z; Investigation: L.Z, H.M, G.S; Resources: Z.J; Data curation: Z.J; Writing – original draft: L.Z, Z.J; Writing – review & editing: Z.J; Supervision: Z.J, A.X; Project administration: Z.J; Funding acquisition: Z.J.

## Funding

This work was supported by the NIH Eunice Kennedy Shriver National Institute of Child Health and Human Development (R01HD102533) and USDA National Institute of Food and Agriculture (2019-67016-29863).

## Data availability

The raw FASTQ files and normalized read accounts per gene are available at Gene Expression Omnibus (GEO) (https://www.ncbi.nlm.nih.gov/geo/) under the accession number GSE272473.

## Supplementary Information

Figure S1. The transcriptional and translational abundances of METTL7A in bovine oocytes and pre-implantation embryo development.

Figure S2. A. An illustrate image showing that ICM is isolated from blastocyst by blastomere biopsy. B. The confirmation of TE and ICM isolation by immunostaining analysis of SOX2 and CDX2 markers, respectively, scale bar = 50µm.

Figure S3. Original images of complete membrane of western blot analysis of SHDA in the METTL7A OE embryos. A. First membrane for SDHA and ACTB. B. Second membrane for ACTB. C. Second membrane for SDHA. OE, METTL7A OE (n = 6); KO, METTL7A knockout (n = 2, note: METTL7A KO experiment was excluded from revised manuscript suggested by the reviewers); VC, vehicle control (n = 7); M, marker (Fisher Scientific cat# 26619). Red arrows indicated SDHA (70 kDa); green arrows, ACTB (45 kDa). Blue arrow, GAPDH (37 kDa). ACTB bands in C were due to insufficient stripping before staining SDHA, but only ACTB bands in B were used for quantification. Original images of complete membrane of western blot analysis of γH2A.X. A. Chemi-plot and overlay-plot of ACTB (45 kDA). B. Chemi-plot and overlay-plot of γH2A.X (17 kDA). OE, METTL7A OE (n = 3); VC, vehicle control (n = 4); M, marker (Fisher Scientific cat# 26619). Due to the long exposure time required, γH2A.X bands did not show up in the overlay-plot.

Table S1. Oligos and antibodies used in this study.

Table S2. Differentially expressed genes in METTL7A OE embryos compared to the control at the 2-cell stage.

Table S3. Differentially expressed genes in METTL7A OE embryos compared to the control at the 8-cell stage.

Table S4. Differentially expressed genes in METTL7A OE embryos compared to the control at the ICM stage.

Table S5. Differentially expressed genes in METTL7A OE embryos compared to the control at the TE stage.

