## Supplementary figures and images for "METTL7A improves bovine IVF embryo competence by attenuating oxidative stress"

### Figure S1

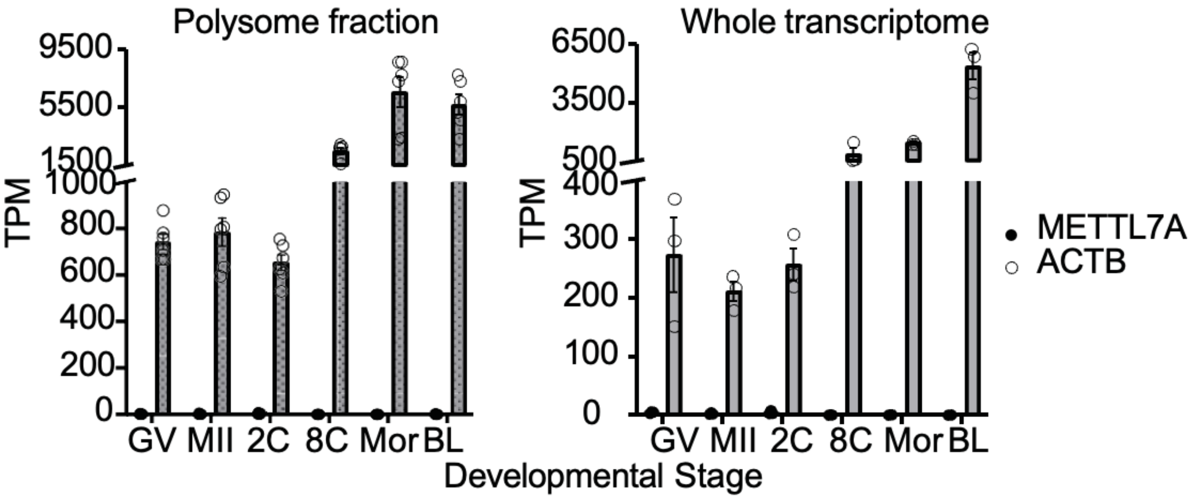

### Figure S2

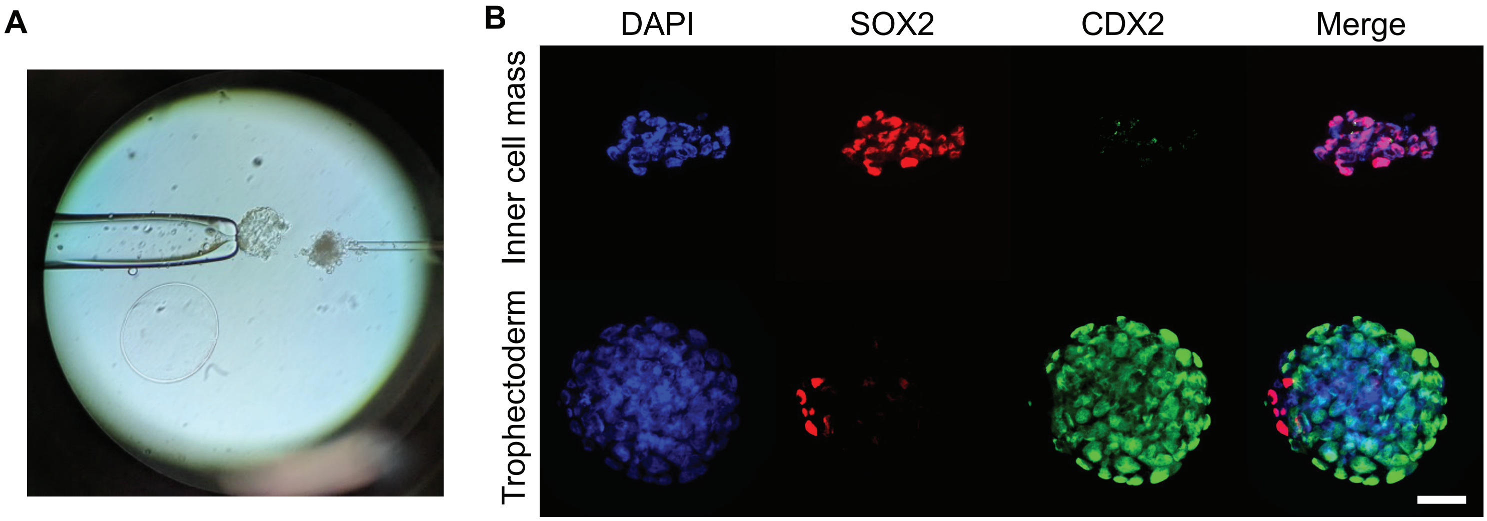

### Figure S3

#### Western blot analysis of SHDA:

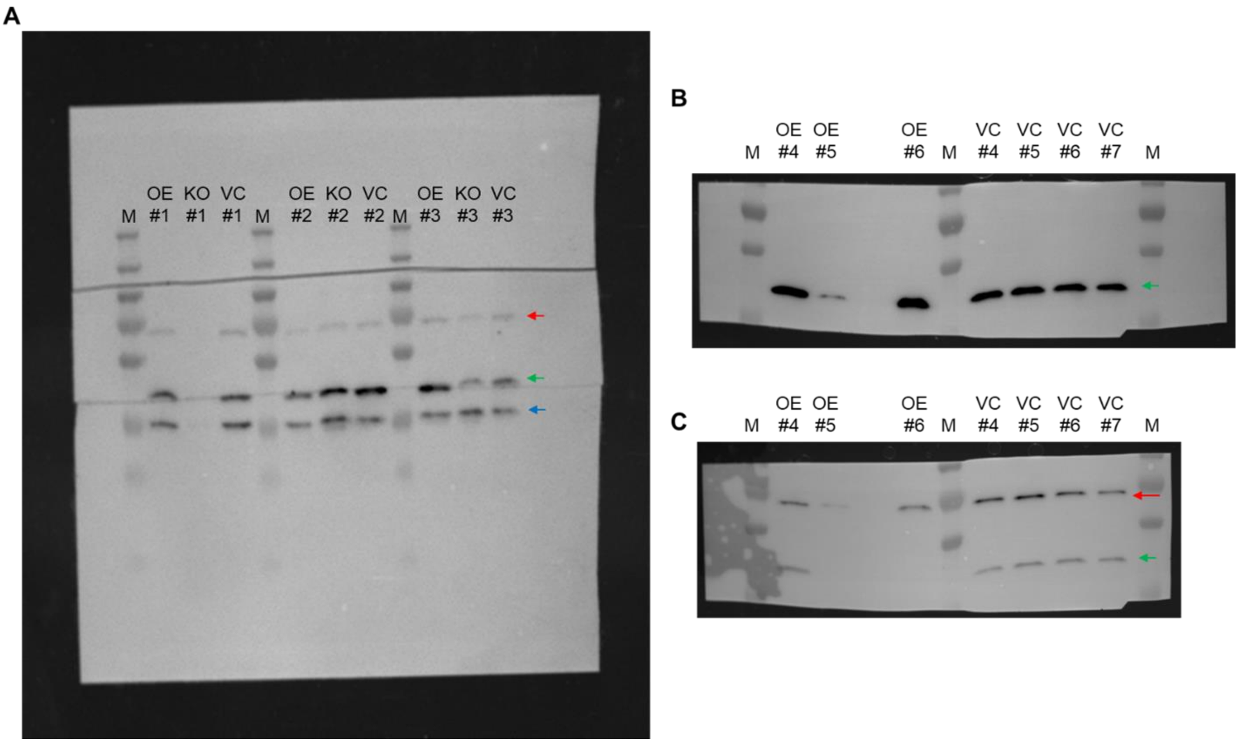


#### Western blot analysis of γH2A.X

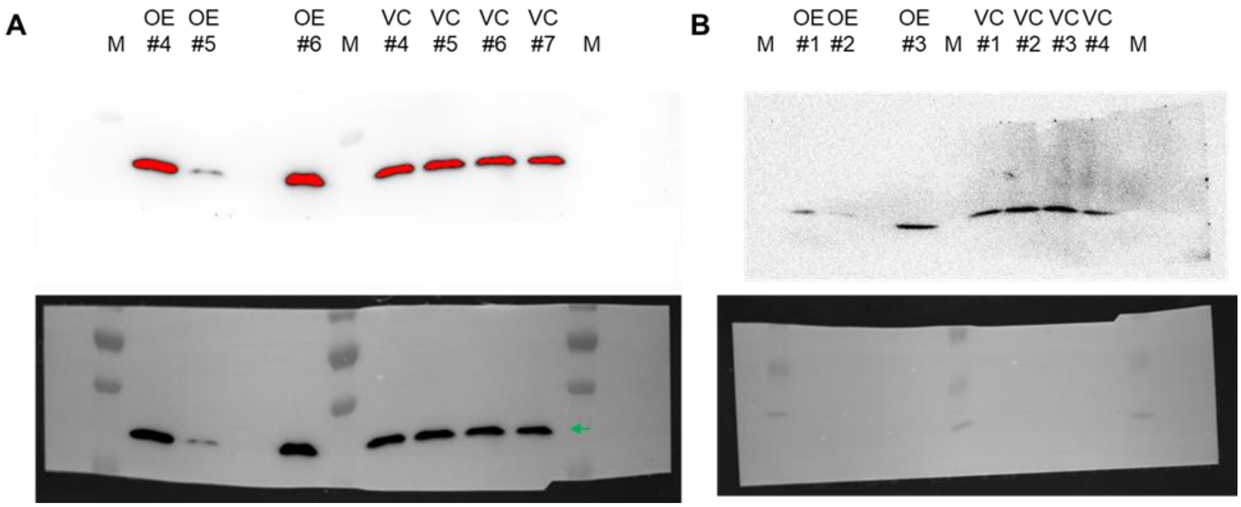
